# WASCO: A Wasserstein-based statistical tool to compare conformational ensembles of intrinsically disordered proteins

**DOI:** 10.1101/2022.12.01.518687

**Authors:** Javier González-Delgado, Amin Sagar, Christophe Zanon, Kresten Lindorff-Larsen, Pau Bernadó, Pierre Neuvial, Juan Cortés

## Abstract

The structural investigation of intrinsically disordered proteins (IDPs) requires ensemble models describing the diversity of the conformational states of the molecule. Due to their probabilistic nature, there is a need for new paradigms that understand and treat IDPs from a purely statistical point of view, considering their conformational ensembles as well-defined probability distributions. In this work, we define a conformational ensemble as an ordered set of probability distributions and provide a suitable metric to detect differences between two given ensembles at the residue level, both locally and globally. The underlying geometry of the conformational space is properly integrated, being one ensemble characterized by a set of probability distributions supported on the three-dimensional Euclidean space (for global-scale comparisons) and on the two-dimensional flat torus (for local-scale comparisons). The inherent uncertainty of the data is also taken into account to provide finer estimations of the differences between ensembles. Additionally, an overall distance between ensembles is defined from the differences at the residue level. We illustrate the interest of the approach with several examples of applications for the comparison of conformational ensembles: (*i*) produced from molecular dynamics (MD) simulations using different force fields, and (*ii*) before and after refinement with experimental data. We also show the usefulness of the method to assess the convergence of MD simulations. The numerical tool has been implemented in Python through easy-to-use Jupyter Notebooks available at https://gitlab.laas.fr/moma/WASCO.

## 1 Introduction

The comparison of protein structures is a crucial problem in structural biology. In the early works [1, 2], the use of root-mean-square deviation (RMSD) was introduced and discussed as a metric between conformations of folded proteins, being later extended to its ensemble version [3]. More recently, Lindorff-Larsen and Ferkinghoff-Borg [4] defined three metrics that allow overall comparison between ensembles of ordered/structured systems, with stronger mathematical guarantees, but using RMSD as a distance between individual conformations, which complicates its extension to disordered structures. Cazals *et al*. [5] used a graph-based representation of the conformational space based on a set of low-energy conformations (i.e. local minima of the potential energy landscape) and compared them with the more suitable Wasserstein distance. To do so, they used the least-RMSD as ground metric between conformations. The methods presented in [4] and [5] are well suited to examine conformational ensembles of molecules that present a well-characterized energy landscape. However, their application to molecules with energy landscapes where low-energy conformations are difficult to identify, as it is the case of IDPs, is inappropriate.

A few recent works have dealt with the comparison of conformational ensembles of IDPs. Huihui and Ghosh [6] focused on averaged conformational properties over ensembles as informative descriptors of their function. They proposed a sequence-decoration metric that classifies IDPs using only their primary structure together with their charge configuration. The same idea of comparing average descriptors was applied by Lazar *et al*. [7], who proposed an ensemble comparison tool based on differences between average pairwise distances. Due to the huge conformational variability of IDPs, it is, however, important to take into account both the average properties as well as the distribution around those averages. Describing IDP conformations as being drawn from probability distributions determining their structure may yield to an important loss of information (or even misleading results) if the whole distribution is reduced to its mean. Even when comparing two (possibly multivariate) Gaussian distributions, the difference between the two depends both on the means and variances [8, 9]; thus, methods for comparing ensembles should ideally include also higher order moments of the probability distributions. This is why a statistical approach that integrates the entire probability law defining an ensemble is crucial to correctly capture the existing differences between disordered ensembles.

In this work, we define a set of probability distributions that characterize at local and global level the highly variable conformations in an ensemble of disordered proteins, and to which we can have access in practice. These probability laws can then be compared using a suitable metric that presents strong mathematical and geometrical guarantees, allowing the identification of residue-specific and overall discrepancies. We also propose an approach to integrate the intrinsic uncertainty of the data within the metric, which enables a more clear identification of the relevant differences between the ensembles. The method has been implemented inside a purely non-parametric framework, avoiding model assumptions, dimensionality reduction or further simplifications that may yield significant loss of information.

In the following sections, we provide an overall description of the proposed methodology, which is exhaustively detailed in the Supplementary Information, together with several cases of applications that illustrate how our method identifies residue-specific and overall discrepancies between conformational ensembles of IDPs or flexible peptides generated for example by molecular dynamics simulations or stochastic sampling techniques.

## 2 Methods

Due to the intrinsic probabilistic nature of IDPs, descriptors of their conformational ensembles should be conceived from a purely statistical point of view. To do so, we seek to locally and globally describe conformational ensembles using well-defined probability distributions and to develop statistical tools allowing their comparison. The main questions to answer are therefore: (1) which is the best way to define those probability distributions? and (2) how these distributions have to be compared to provide quantitative information about similarities and differences between ensembles?

### 2.1 Defining conformational ensembles as a set of probability distributions

IDP ensembles can be described at both local and global scales, providing complementary information. We aim at defining an ordered set of probability distributions that account for the highly variable structure of the ensemble and, above all, that can be estimated in practice from a set of sampled conformations.

The most important aspects of the local structure can be described by the dihedral angles (*φ, ψ*) for each amino acid residue along the sequence. Therefore, for each residue, the ensemble is locally characterized by a two-dimensional random variable (*φ, ψ*) or, in other words, by a probability distribution supported on the two-dimensional flat torus 𝕋^2^ [10,11]. If we denote such distribution as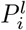, for the residue at the *i*-th position, we define the local structural descriptor of an ensemble as the *L*-tuple

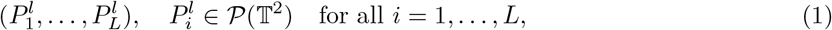

where *L* is the sequence length and 𝒫 (𝕋^2^) denotes the space of probability distributions supported on𝕋^2^.

Describing the global structure is a less trivial task. The use the absolute positions of the atoms and an absolute reference frame for the entire ensemble is not an appropriate description as it is sensitive to rigid-body motions. Therefore, our approach uses the relative positions of all pairs of residues along the sequence, which are invariant under rigid-body motion. More precisely, we define the position of a given residue as the the position of its *C*_*β*_ atom when it exists and of its *C*_*α*_ atom otherwise. If *i, j* ∈ *{*1, …, *L}, i* ≠ *j*, denote two different sequence positions, let 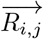 be the three-dimensional random variable determining the relative position of *j*-th residue with respect to the *i*-th one. If we denote 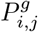 the probability distribution associated to 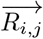, we define the global structural descriptor of an ensemble as the (*L*(*L* − 1)*/*2)-tuple

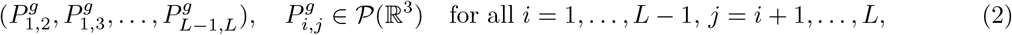

where *L* is the sequence length and 𝒫 (ℝ^3^) denotes the space of probability distributions supported on the three-dimensional euclidean space.

### 2.2 Accessing empirical probability distributions from sampled conformations

Estimating the local structural descriptor (1) is immediate as we have direct access to dihedral angles (*φ, ψ*) from the sample of conformations. Therefore, the local structural descriptor will be estimated by its *empirical* counterpart

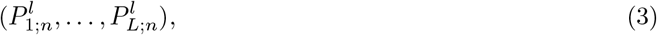

where each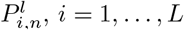, is the empirical probability distribution of 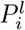, and *n* is the number of conformations constituting the sample. Such empirical probability distributions are commonly represented through Ramachandran maps [12].

Obtaining a sample of 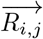 from the set of conformations is less direct. To compute a set of comparable 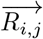 vectors from all conformations, their coordinates must be expressed on the same reference system.

To do so, we first define a reference frame at the *i*-th residue, using only the positions of the *i*-th *C*^*′*^, *C*_*α*_ and *N* ^*H*^ atoms. This frame, whose construction is detailed in the SI, is a meaningful representation of the spatial pose of each residue.

The reference frame associated to each residue *i* ∈ *{*1, …, *L}* allows to express the relative positions of all residues *j* ≠ *i* with respect to *i*. Moreover, the definition of a reference system allows the *superposition* of all the conformations in the ensemble. This is illustrated in Figure 1, for three conformations. Consequently, for every *j* ≠ *i*, we will have access to *n* realizations of the random variable 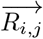 or, in other words, to a point cloud in the three-dimensional Euclidean space, representing a sample drawn from the distribution of 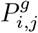. Therefore, the global structural descriptor of the ensemble (2) will be estimated by its *empirical* counterpart

**Figure 1:**
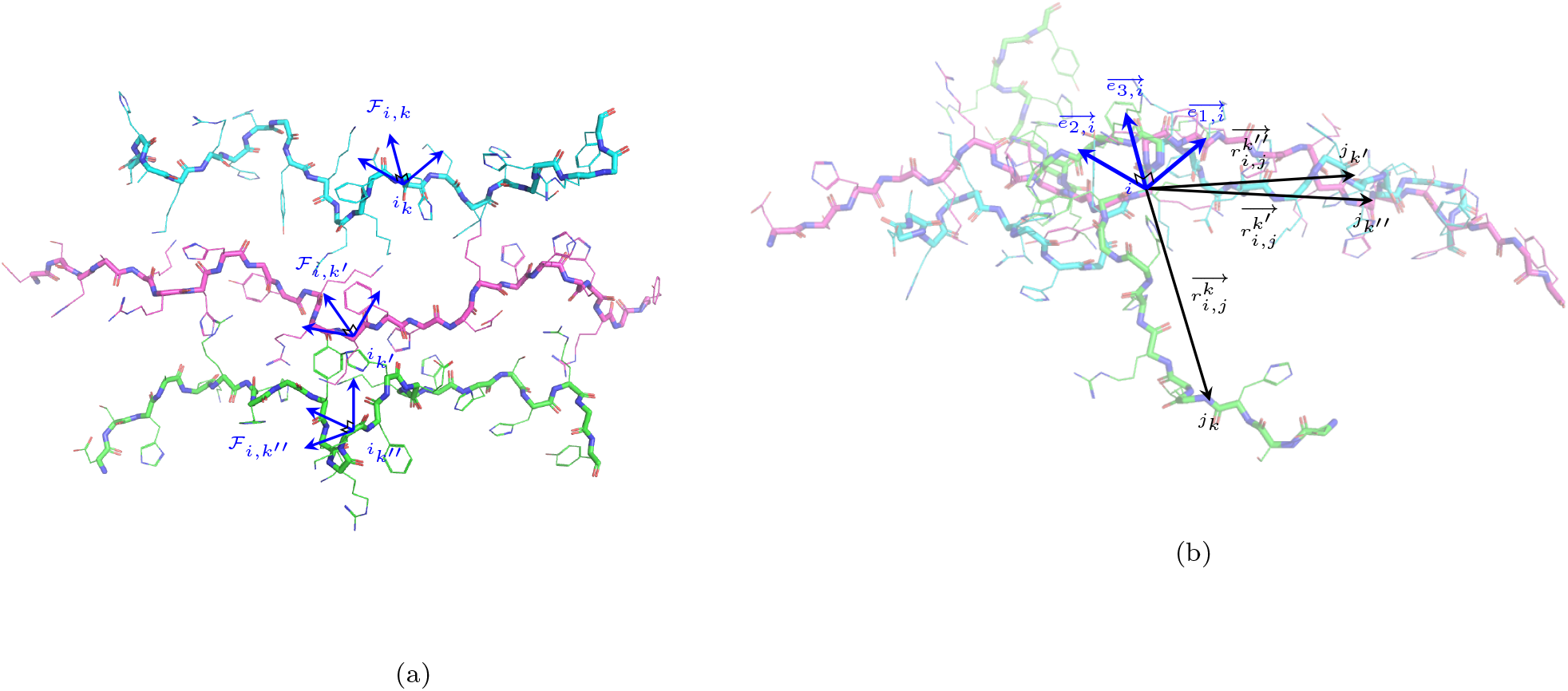
Illustration of how samples of global structural descriptors are obtained, for a pair of positions *i, j* along the sequence. In (a), the reference frame is built for every conformation at residue *i*. In (b), all the frames are superimposed using this reference frame. Then, for any *j* ≠ *i*, the vectors 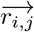 constitute a sample of 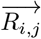.

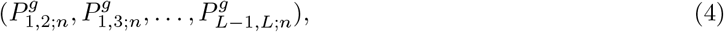

where 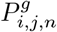 is the empirical probability distribution of 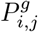, for all *i* = 1, …, *L* − 1, *j* = *i* + 1, …, *L*. An example of a pair of samples of 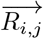 is presented in Figure S4.

### 2.3 Distances between local and global structural descriptors

After defining the local and global structural descriptors of an ensemble as an ordered set of probability distributions, the choice of a suitable metric allowing inter-ensemble comparisons becomes the subsequent question to deal with. The basic properties that such a metric should have are:

1. Satisfying the mathematical properties that define a distance (i.e. being 0 if an only if the two compared distributions are identical, symmetry and triangle inequality),
2. Integrating the geometry of the underlying space.

The use of metrics between probability distributions is not new in structural biology. For instance, Ting *et al*. [13] used Hellinger distance to detect differences between (*φ, ψ*) distributions. However, this metric does not take into account the geometry of the underlying space (in particular here, its periodicity). A symmetrized Kullback-Leibler (KL) or the Jensen-Shannon (JS) divergence was used in [4, 14] to compare ensembles of ordered systems. This metric has a firm interpretation, based on information theory (in particular the JS divergence is the square of a metric). However, it still misses the geometrical reliability and does not satisfy triangle inequality, which makes comparisons between multiple ensembles difficult to interpret.

Besides satisfying conditions 1 and 2, the Wasserstein distance, derived from the theory of Optimal Transport (OT), provides both strong theoretical guarantees [15] and attractive empirical performance [16]. Moreover, the Wasserstein distance (also known as the ‘earth mover distance’) has a physical interpretation, as it is defined as the minimum transportation cost needed to reconfigure the mass of one probability distribution to recover the other. For an introduction to this Theory, we refer to [16]. Most of the applications of OT are related to the very active field of machine learning, notably in the framework of generative networks [17], robustness [18] or fairness [19], among others. With some notable exemptions [5, 20–23], Wasserstein distance has not been widely used in structural biology. More related to our work, in [5], Cazals *et al*. used Wasserstein distance to compare energy landscapes sampled from conformational ensembles. Recently, it was used in [23] to define statistical tests assessing differences between (*ϕ, ψ*) distributions. The incorporation of the underlying geometry to its definition makes it a well-adapted metric to measure distances between local and global structural descriptors of the ensembles. Details and important considerations regarding its practical computation are given in the SI.

### 2.4 The comparison tool

Consider two ensembles *A, B*, associated to two protein sequences of equal length *L*, and let *n*_*A*_, *n*_*B*_ be their number of conformations, respectively. We define the differences between local structural descriptors of *A* and *B* as the *L*-tuple of Wasserstein distances

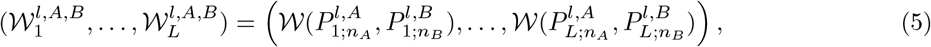

where 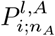 (resp. 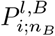) denotes the *i*-th distribution of the empirical local structural descriptor (3) of ensemble *A* (resp. *B*). Statistical tests to assess whether any 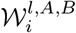 is significantly different from zero have been recently defined in [23]. The second of the introduced techniques is better adapted to our problem, as it only detects the more important discrepancies and accepts slight differences that may arise from experimental or computational procedures. This is discussed in detail in [23]. Consequently, together with the *L*-tuple (5) of distances comparing local structural descriptors, we are able to supply a *L*-tuple of *p*-values accounting for the statistical significance of the corresponding distances:

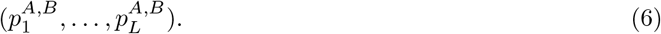

Recall that a small *p*-value 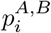 indicates strong evidence that the *true* distance that 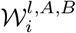 estimates is different from zero. In other words, small *p*-values show significant differences between the corresponding local structural descriptors. Therefore, the vector (6) enables the identification of those residues where the differences are more important, and those residues for which differences can be assigned as non-significant.

Analogously, the difference between global structural descriptors of *A* and *B* is defined as the (*L*(*L* − 1)*/*2)tuple

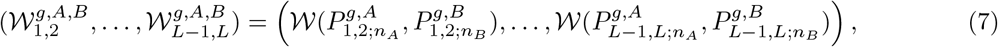

where 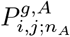 (resp. 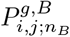) denotes the *i, j* distribution of the empirical global structural descriptor (4) of ensemble *A* (resp. *B*). Note that (7) can be more naturally represented as a triangular (*L*− 1) *×* (*L*− 1) matrix *W*^*g,A,B*^, whose elements are given by 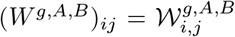. Graphically, the matrix *W* ^*g,A,B*^ is represented using a color scale to fill the coefficients according to distance values. As the diagonal will remain empty, it will be filled with the local distances (5). This will also allow to assess whether changes on local structural descriptors are related with changes in global structural descriptors and to compare both scales within the same representation.

#### 2.4.1 Accounting for uncertainty

The variability in experimental and simulated structures causes uncertainties and statistical noise that may significantly bias the distance estimation. For example, when running a MD simulation, independent replicas of the same simulation setup may results in non-negligible differences that distort the analysis of the comparison matrices. The same may occur when comparing two uniformly chosen subsets of conformations from an ensemble generated by stochastic sampling techniques [24, 25]. In order to soften the effect of uncertainty and to obtain *net* estimates of the differences between a pair of ensembles, we will use (if available) independent replicas of the same ensemble. These replicas may also be produced by uniform subsampling of the set of conformations. However, special care must be taken when subsampling MD trajectories as the convergence of the simulation must be ensured for the subsamples to be representative of the entire ensemble.

Let 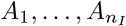 (resp.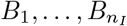) be *n*_*I*_ independent replicas of ensemble *A* (resp. B). The corrected difference between local structural descriptors of *A* and *B* is defined as the *L*-tuple

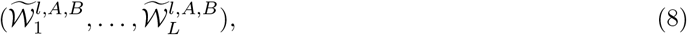

where each corrected distance is defined as

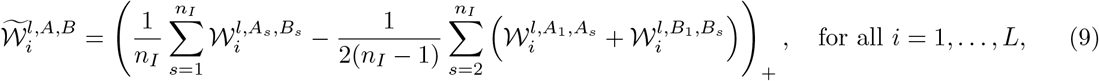

where, for any real number *x*, (*x*)_+_ = *x* if *x >* 0 and (*x*)_+_ = 0 otherwise. The first term in (9) is an average of *n*_*I*_ Wasserstein distances between *n*_*I*_ paired independent replicas of *A* and *B*. As it was shown in [26], an average of Wasserstein distances between sub-samples of the same population is a pertinent estimate of the Wasserstein distance between the two entire populations that, in addition, conserves the properties that mathematically define a distance. Therefore, this first term estimates the Wasserstein distance between the entire populations of *A* and *B* (conceived as the union of all independent replicas), softening the variability. To this *brutto* inter-ensemble difference, we subtract an average of the Wasserstein distances between independent replicas of the same population (intra-ensemble). Note that, for the sake of computational simplicity, we just compared the first replica of each ensemble with the subsequent ones. Of course, distances between all pairs of replicas can be added to this term. The same applies if *n*_*I*_ is different for *A* and *B*; both terms in (9) can be accordingly adapted. As it is illustrated in Section 3, the use of corrected distances (9) contribute to reduce the noise coming from structural uncertainty and help to emphasize residue-specific differences in the matrix representation. For the distances between global structural descriptors, the correction is performed analogously.

#### 2.4.2 Setting an interpretable scale

When defining an absolute distance or score between conformational ensembles, providing the clues to ease its interpretation is crucial. The problem of interpreting unbounded metrics with no intrinsic reference values has been widely discussed since the introduction of RMSD for the comparison of pairs of conformations [1, 2]. Here, we do not seek to define any cutoff to binarize the resulting matrices, but to provide a more informative continuous scale. To do so, we aim at quantifying the magnitude of the inter-ensemble distances compared to the intra-ensemble ones, using the uncertainty estimate as a reference. If we denote as 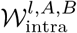(resp.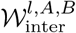) the first (resp. second) term in (9), the score

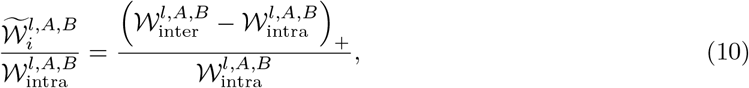

reflects in which proportion the inter-ensemble distances exceed the “default” intra-ensemble ones. Once again, this score is analogously defined for differences between global structural descriptors.

#### 2.4.3 An overall distance between ensembles

In some situations, it may be of interest to perform overall comparisons between multiple ensembles. To do so, moving from a residue-specific analysis to a comparison at the whole structure level might be preferable. The definition of a score for the overall ensemble has been addressed for ordered systems [4]. Here, we propose to define such a score by aggregating all the residue-specific distances computed using the above-described methods. We recall that if *d*_1_, …, *d*_*L*_ are *L* distances defined on *L* metric spaces 𝒳_1_, …, 𝒳_*L*_, the function 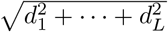 is a distance on the product space 𝒳_1_ *×* … *×*𝒳_*L*_. Consequently,

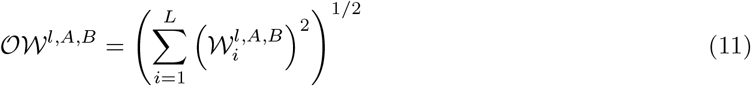

is a distance on the product space of all dihedral angles along the sequence and, therefore, serves to quantify the *overall local discrepancy* between a pair of ensembles. Analogously,

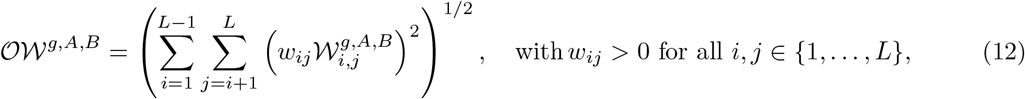

is a distance on the product space of all pairwise relative positions of the residues in both ensembles, and serves to quantify the *overall global discrepancy*. Note that we have assigned a positive weight *w*_*ij*_ to each global distance in (12). This allows to consider distances between specific residue pairs as more relevant than the others when computing the overall discrepancy [27]. For instance, we can highlight differences between global structural descriptors that appear for residue pairs that are far from each other in the sequence, i.e. large |*i*− *j*|, and neglect distances between neighboring residue pairs, i.e small |*i*− *j*|. This can be done by choosing *w*_*ij*_ as an appropriate increasing function of |*i* − *j*|, as

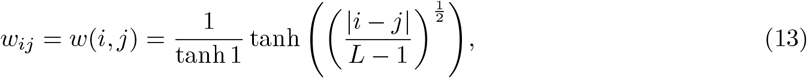

which satisfies *w*_*i,i*_ = 0 for all *i* and *w*_1,*L*_ = *w*_*L*,1_ = 1.

The drawback of this definition of the overall distance is that it does not take into account the uncertainty discussed in Section 2.4.1. To solve this problem, the same strategy to define a global score can be performed by replacing each 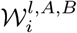(resp.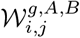) by its corresponding corrected distance 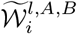 (resp.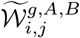) in (11) (resp. (12)). However, this strategy makes the triangle inequality for the overall metric no longer satisfied. Both scores can be implemented by the practitioner and used depending on the specific comparison context.

### 2.5 The Jupyter Notebook

The WASCO comparison tool has been implemented through an easy-to-use Jupyter Notebook. It is available at https://gitlab.laas.fr/moma/WASCO, together with its installation guidelines and detailed implementation instructions. The notebook takes a pair of ensembles as input and returns the comparison results through the matrix defined in Section 2.4, containing global and local differences. The user can choose to correct the computed distances by uncertainty, as proposed in Section 2.4.1. When independent replicas are not provided as input, subsampling is used to emulate them. If this correction is performed, results are displayed in the interpretable scale defined in Section 2.4.2. The overall score defined in Section 2.4.3, aggregating the corrected distances, is also returned by the tool.

Ensembles can be provided as input in several of the most common data formats. WASCO accepts one .xtc file per replica, together with a .pdb file including the topology information of the molecule, one multiframe .pdb file per replica or a folder per replica containing one .pdb file per conformation. The user can also choose to compare ensembles for sequence segments (of equal length) instead of the entire sequence. Details are provided in the notebook documentation.

Due to the large number of Wasserstein distances to be computed (*L*(*L* − 1)*/*2 + *L* per pair of replicas), the computation time might be considerably high. The number of conformations constituting the ensemble also has a significant impact, due to computational limitations of the existing OT algorithms when sample sizes and dimension increase. In order to return results within a reasonable amount of time, WASCO computes Wasserstein distances in parallel. The required CPU time depends on the number of conformations, replicas and sequence length of the ensembles. For small proteins of *L* ∼ 30 and ensembles or reasonable size *n*_*A*_, *n*_*B*_ ∼ 10^4^, the CPU time using 20 threads is less than 15 minutes using a standard computing server. However, for larger proteins of *L* ∼ 150 and large ensembles with *n*_*A*_, *n*_*B*_ ∼ 10^5^, the CPU time using 20 threads goes up to some hours. Additionally, comparing large ensembles of substantially longer sequences (*L* ≫ 150) might cause memory problems, as all pairwise relative positions for every conformation need to be stocked. Therefore, the suitability of the sizes of the ensembles must be considered before launching WASCO. Adapting WASCO to longer sequences with large conformational ensembles remains an objective for future work.

## 3 Results

In this section, we present several applications to illustrate the different possibilities enabled by WASCO. In all the cases, the distances between local and global structural descriptors were corrected for uncertainty using (9), as independent replicas were available. Both overall local and global discrepancies between pairs of ensembles were computed plugging the corrected distances in (11) and (12), as discussed in Section 2.4.3. The weight function (13) was used to highlight differences between residue pairs far from each other in the sequence and reduce differences between neighboring amino acids. Note that this weighting is considered only to compute the overall distance (12), and not to depict distance values in the matrix representation, which correspond to the interpretable scale defined in Section 2.4.2.

### 3.1 Comparison of ensembles produced by MD simulations using different force-fields

We applied WASCO to compare the results of MD simulations using different force-fields presented in [28] for two flexible peptides showing a significant propensity to form poly-l-proline type II (PPII) structures. Four different force-fields, having demonstrated relatively good performances to simulate IDPs were applied: AMBER ff99SB-disp, AMBER ff99SB-ILDN, CHARMM36IDPFF, and CHARMM36m (details and references to these force-fields can be found in [28]). For simplicity, we will refer to these force-fields as disp, ildn, c36idp and c36m, respectively. As independent replicas for each simulation were available, we could perform the correction for uncertainty (9).

Figure 2 presents the output of WASCO for several pairwise comparisons of conformational ensembles of Histatin-5 (Hst5) obtained with the different force-fields. The matrices and the overall dissimilarities suggest that the generated structures are closer (in Wasserstein distance) when they are simulated using c36idp and c36m (which we can define as group-I), or disp and ildn (group-II). This is not surprising as group-I are versions of CHARMM and group-II are versions of AMBER. Indeed, matrices (a) and (b), comparing force fields inside group-I and inside group-II respectively, present overall global differences which are small compared to those of panels (c) and (d), which compare force-fields of different groups. The same conclusion can be extracted by comparing the magnitude of the scales of both pairs of matrices. The two remaining comparisons (ildn vs. ildn and c36m vs. disp) are not included in Figure 2 as the corresponding matrices are qualitatively equivalent to (c) and (d). We note that similar observations have been made when comparing ensembles of folded proteins generated using related force-fields [14, 29].

**Figure 2:**
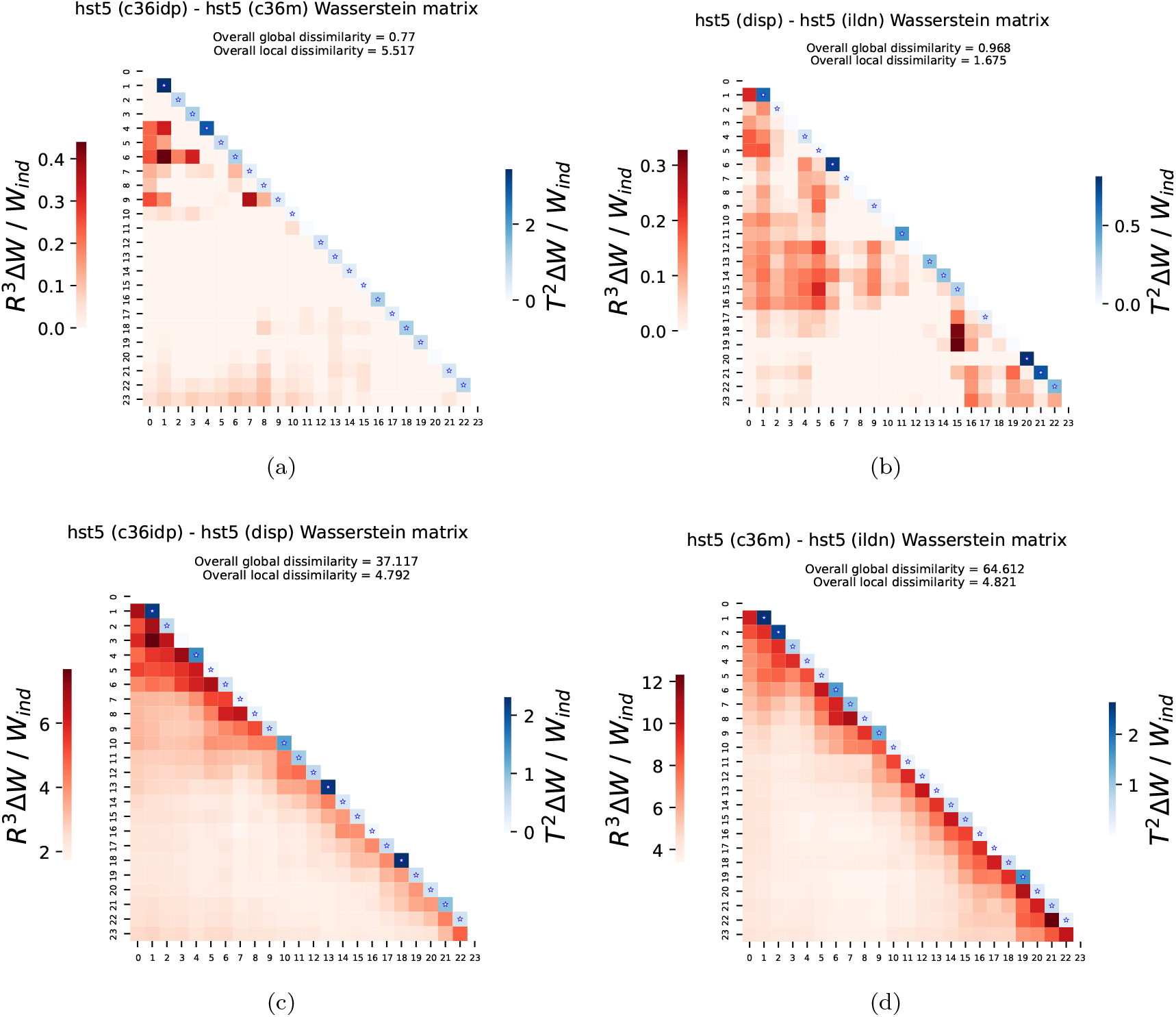
Comparison of Molecular Dynamics simulations of Hst5 ensemble using different force-fields. The color scale indicates the proportion of intra-ensemble differences that corresponds to the observed corrected inter-ensemble differences, as defined in (10). The coefficients in the lower-triangle (in red) correspond to the global differences. The coefficients along the diagonal (in blue) correspond to the local differences. Blue stars indicate that the corresponding local corrected distance is significantly different from zero (the associated *p*-value (6) is smaller than *α* = 0.05). Note the very different scales used in the different plots.

Matrices returned by WASCO also allow a residue-specific analysis of the distances. In Figure 2, panels (c) and (d) show that the most relevant global differences appear in regions close to the diagonal (i.e. between residue pairs close in the sequence), where the inter-ensemble corrected distances rise up to 6-7 times the intra-ensemble ones. This is not the case when comparing force-fields inside the same group, as the largest differences appear in more internal matrix regions (i.e. between residue pairs more distant in the sequence). However, these corrected differences represent less than the half of the intra-ensemble distances. The information displayed on the diagonal allows the detection of the residues where the local conformation change more abruptly between force-fields. These local changes are more localized, contrary to the observed behaviour of global differences, which appeared for regions inside the lower triangle and not for isolated pairs of amino acids. In some cases, substantial local distances appear in residues where global structure also changes (see, for example, residues next to the N-terminus in (a,c)). However this correspondence is not observed in all matrices.

We repeated the same analysis for MD simulations of PEP3 with the same force-fields. Results are presented in Figure 3. Here, the discrimination between the two force-field families is not observed. Nonetheless, we still observe that structures simulated with disp and ildn are very close in Wasserstein distance (Figure 3b). Indeed, the overall global dissimilarity is substantially smaller than these of the remaining comparisons. Only inter-ensemble corrected differences representing about the 20% of the intra-ensemble ones appear for residues at the C-terminus. The distances between c36idp and c36m are now higher than for Hst5, and corrected differences of the same magnitude than the intra-ensemble ones appear in the interior of the matrix. The same behavior when comparing force-fields of different groups appear for PEP3. See, for instance, that substantial differences arise between relative positions of residues at opposite terminus in panel (d), which are highly weighted when computing the overall global discrepancy. One intriguing observation is that while there are substantial differences between disp and ildn (and between c36idp and c36m), simulations with c36idp and c36m used the same water model (the CHARMM-modified TIP3P water model) and the disp and ildn simulations also used very similar water models (TIP4P-D and a slightly variant of this) [28]. Overall, these results are complementary to those presented in [28], which mainly focused on secondary structure differences among ensembles, and they show the ability of WASCO to identify differences at both local and global scales.

**Figure 3:**
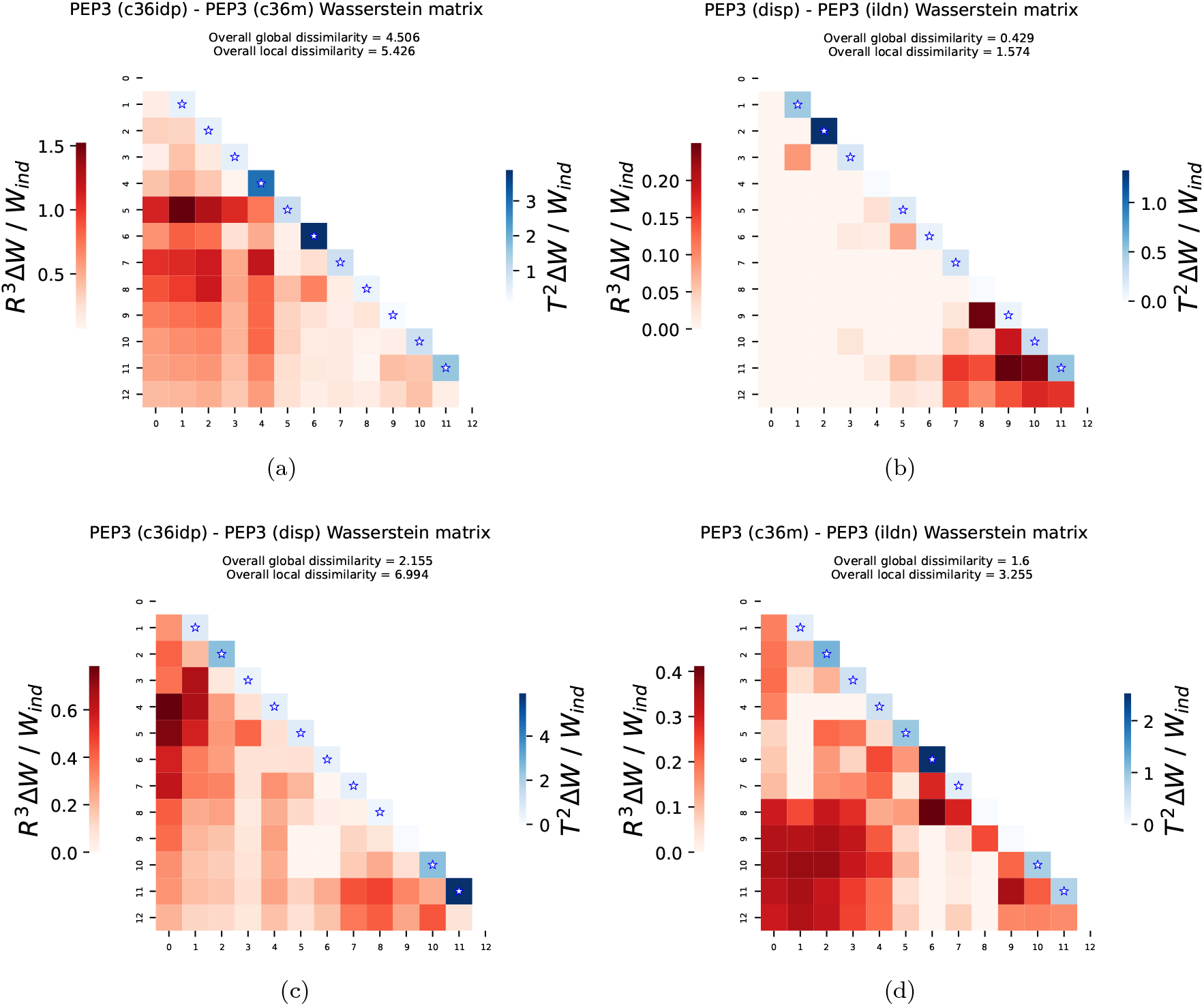
Comparison of Molecular Dynamics simulations of PEP3 ensemble using different force fields. The color scale indicates the proportion of intra-ensemble differences that corresponds to the observed corrected inter-ensemble differences, as defined in (10). The coefficients in the lower-triangle (in red) correspond to the global differences. The coefficients along the diagonal (in blue) correspond to the local differences. Blue stars indicate that the corresponding local corrected distance is significantly different from zero (the associated *p*-value (6) is smaller than *α* = 0.05).

### 3.2 Online assessment of the convergence of MD simulations

Ensemble comparisons have previously been used to assess convergence in MD simulations of folded proteins [14, 29, 30]. We here propose to use the overall ensemble distances (defined in Section 2.4.3) to examine the convergence of an MD simulation of a disordered protein. Moreover, this can be done on-the-fly to assess whether the simulation can be stopped. Let *T* denote the current simulation time and let 0 *< t*_1_ *< t*_2_ *<* … *< t*_*k*_ = *T* be *k* time points. If we denote *A*_*t*_ the conformational ensemble simulated up to time *t*, we can compute the overall distances

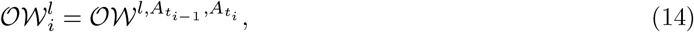

defined in (11), for all *i* = 2, …, *k*. For each *i*, 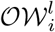 corresponds to the overall local distance between the ensemble from *t* = 0 to *t* = *t*_*i*_ and the ensemble from *t* = 0 to *t* = *t*_*i*−1_. Analogously, we compute the overall global distances

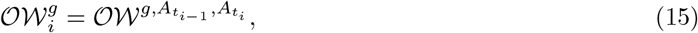

as defined in (12). Then, the representation of 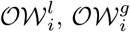 with respect to the *t*_*i*_ indicates whether the simulation has converged or not. Note that the distances 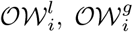 can never be equal to zero, as they are empirical distances which *converge* to zero when the sample size tends to infinity. Therefore, convergence will be assumed when the profile approaches asymptotically to zero. Nevertheless, this criteria provides a stronger evidence of non-convergence, as the achievement of a zero asymptote for (15), even if necessary, may not be sufficient to guarantee convergence. If we resolve that the simulation must keep going until time *T* ^*′*^ *> T*, it suffices to add 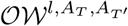 and 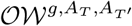 to each curve and recheck.

Figure 4a presents the evolution of the online overall distances for PEP3 simulated with the four force-fields introduced in Section 3.1. We observe that all the curves exhibit an asymptote at zero after the 80% of simulation time and therefore, convergence has been achieved in all cases. This is not the case for the simulation in Figure 4b, corresponding to a 1,000 ns simulation of the K-18 domain of Tau using the AMBER ff99SB*-ILDN force-field and the TIP4P-D water model (Sthitadhi Maiti and Matthias Heyden, unpublished). Here, we clearly observe that no asymptote at zero is present and therefore that the simulation has not attained convergence. This result was expected due to the length of the protein (129 amino acids) and the reduced simulated time.

**Figure 4:**
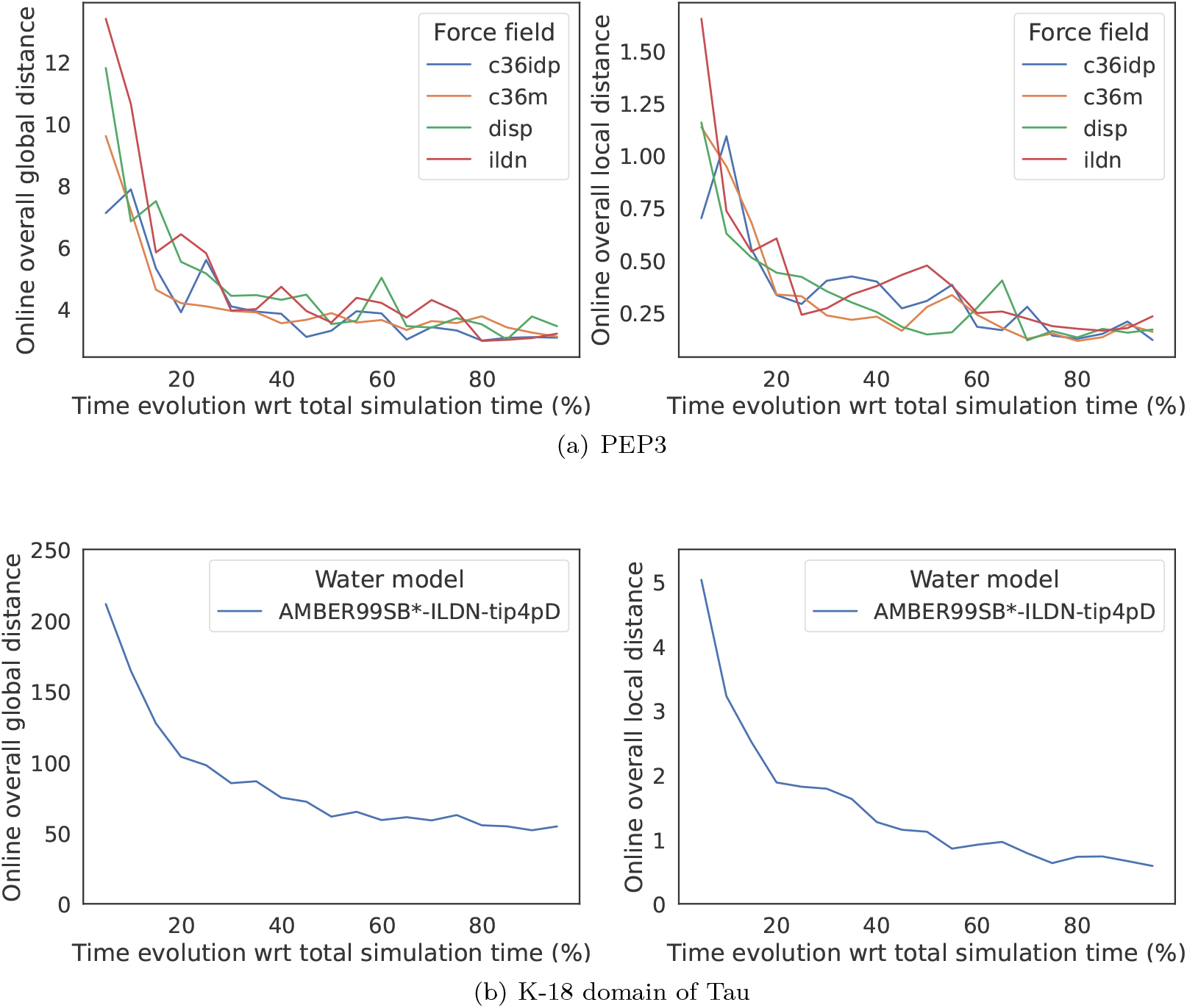
(a) Online convergence analysis for PEP3 ensemble simulated with force-fields c36idp, c36m, disp and ildn. (b) Online convergence analysis for K-18 domain of Tau ensemble simulated with AMBER ff99SB*-ILDN and TIP4P-D water model. In abscissas, the percentage of simulation time, divided in 20 equally spaced time intervals. In ordinates, the overall distances between the ensembles simulated at the extremes of the time intervals. The left (resp. right) column presents the evolution of 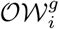(resp. 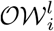) with respect to time.

### 3.3 Structural impact of SAXS ensemble refinement

Using Hst5 as example, we applied WASCO to evaluate the structural impact of SAXS refinement with the Ensemble Optimization Method (EOM) [31] on the resulting ensemble. We first compared the Hst5 ensemble simulated with Flexible-Meccano [24, 32] with the refined one using previously reported SAXS data [33]. The results are presented in Figure 5. Note that a previous EOM analysis of these data suggested that Hst5 in solution is slightly more extended that the random coil model generated with Flexible-Meccano [33]. Small but significant differences were observed at the central part of the peptide (from residues 6 to 13). Most probably, this central region has been conformationally modified during the refinement to account for the overall expansion of the peptide in solution [31]. Moreover, we observed highly significant local distances that propagate towards the interior of the matrix. In other words, these residues with large local distances conformationally influence their closest neighbours. Intriguingly, this propagation seems to only propagate towards the C-terminus.

**Figure 5:**
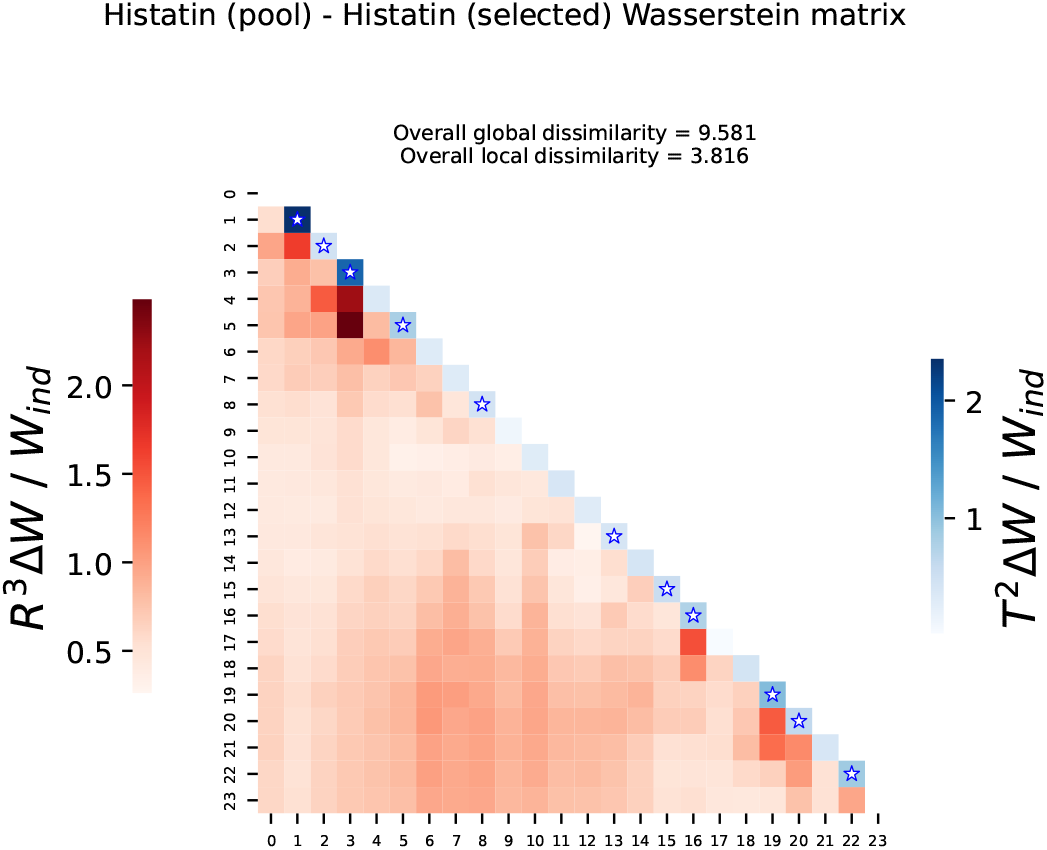
Comparison of Hst5 ensemble before and after filtering with experimental SAXS data. The color scale indicates the proportion of intra-ensemble differences that corresponds to the observed corrected inter-ensemble differences, as defined in (10). The coefficients in the lower triangle (in red) correspond to the global differences. The coefficients along the diagonal (in blue) correspond to the local differences. Blue stars indicate that the corresponding local corrected distance is significantly different from zero (the associated *p*-value (6) is smaller than *α* = 0.05).

We next assessed whether the direction in which conformations are built have a structural effect and change the refined ensemble. To do so, we generated two Hst5 ensembles using a stochastic sampling method similar to Flexible-Meccano but using a different sampling strategy [25], where the chains were built either from N-to-C or from C-to-N. When using these two ensembles to fit the experimental curve, the resulting distance matrices displayed very similar features for local and global distances (Figures 6a and 6b), suggesting that the chain-building direction does not have a relevant effect. In both cases, a systematic increase in the distances is observed for the central residues, as observed in the previous analysis (Figure 5).

**Figure 6:**
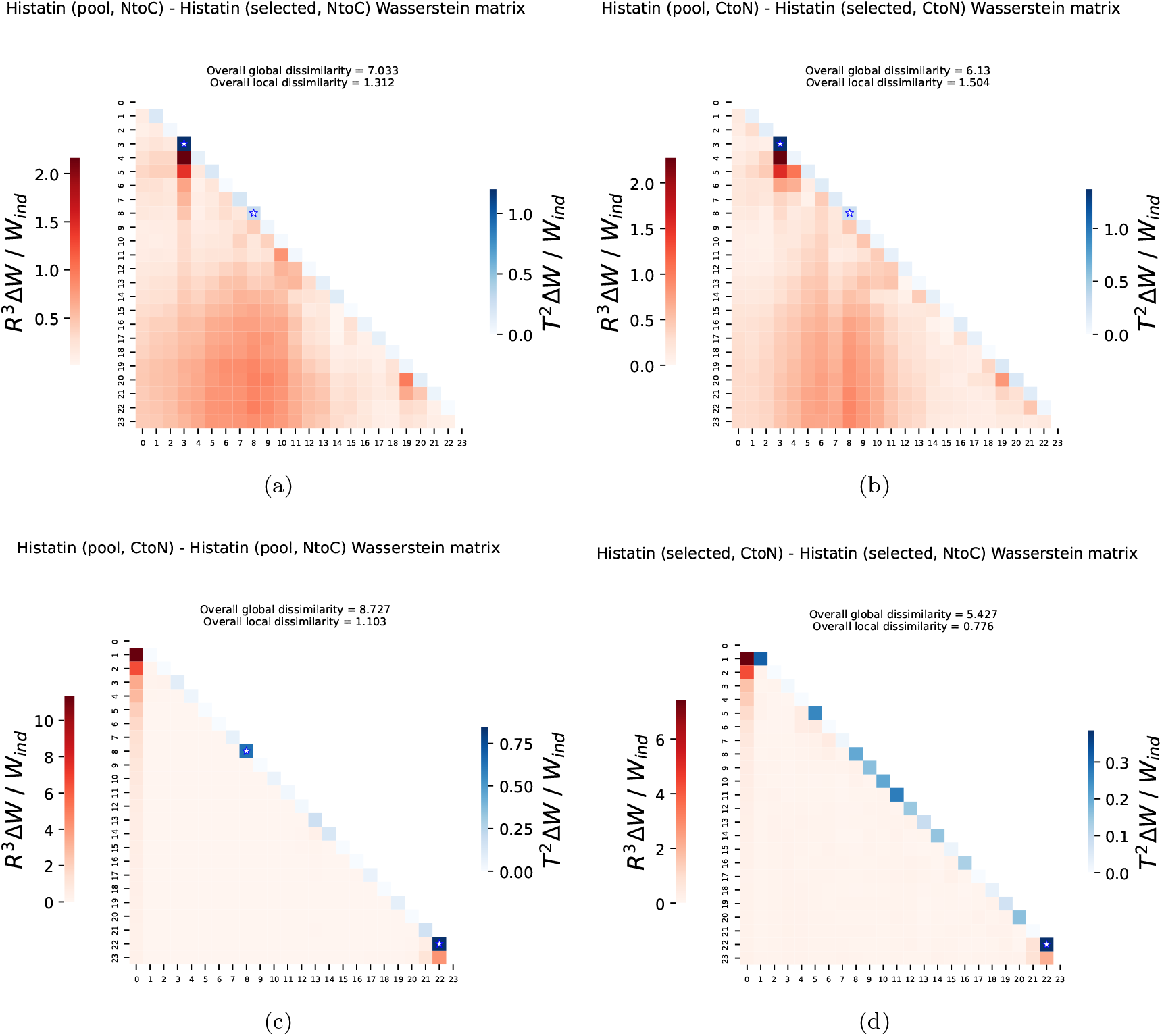
comparison of Hst5 ensembles before and after filtering with experimental SAXS data. The ensemble was simulated from (a) N-to-C or from (b) C-to-N. (c) Comparison of Hst5 ensembles generated from N-to-C and C-to-N. (d) comparison of the N-to-C and C-to-N SAXS refined. In all matrices, the color scale indicates the proportion of intra-ensemble differences that corresponds to the observed corrected inter-ensemble differences, as defined in (10). The coefficients in the lower triangle (in red) corresponds to the global differences. Coefficients along the diagonal (in blue) correspond to the local differences. Blue stars indicate that the corresponding local corrected distance is significantly different from zero (the associated *p*-value (6) is smaller than *α* = 0.05).

In a recent study, ENCORE was used to show that refined ensembles were closer to each other than different input ensembles [14]. This can also be illustrated using WASCO, by comparing the Hst5 ensembles generated in both directions before and after the filtering with SAXS data (Figures 6c and 6d). These comparisons clearly showed that both global and local differences were smaller for the refined ensembles than for the input ones, as observed when comparing the maximum values of the corresponding color scales. As we were comparing very similar ensembles, we expected the distances to be small. Nevertheless, we observe one significant local difference on the diagonal in Figure 6c that disappeared after refinement.

## 4 Discussion

We have presented a novel method to compare conformational ensemble models of highly flexible proteins. WASCO is based on a non-parametric framework: local and global structural descriptors of the conformational space are defined as distributions and do not rely on probabilistic or statistic models. This allows capturing the entire variability of the ensemble without information loss. The distributions are compared using the Wasserstein distance, which has strong mathematical guarantees and respects the geometry of the underlying space. To this metric, we incorporated the structural uncertainty presented in experimental and simulated ensembles. Using this strategy, WASCO highlights the relevant differences between ensembles. We have illustrated several possible applications of WASCO as an additional tool for the investigation of IDPs and flexible peptides. It provides complementary information with respect to other tools to analyze and compare conformational ensembles based on global descriptors, such as the radius of gyration [34] or secondary structure propensities [28]. WASCO has been implemented in an open source Jupyter Notebook. The main drawback of this implementation is its inability to deal with considerably large ensembles. Adapting WASCO to larger chains remains a future work, as well as its extension to compare multi-domain proteins and coarse-grained models.

## Acknowledgements

We are grateful to Francesco Pesce, Sthitadhi Maiti and Matthias Heyden for providing useful data. We thank Gabriella Gerlach, Frederik Emil Thomasen and Philipp Schake for their helpful discussions and valuable feedback on WASCO implementation.

This work has been partially supported by the French National Research Agency (ANR) through grant ANR-19-P3IA-0004, the LabEx CIMI (ANR-11-LABX-0040) and EpiGenMed (ANR-10-LABX-12–01) within the French State Programme “Investissements d’Avenir”, by the European Research Council under the European Union’s H2020 Framework Programme (2014–2020)/ ERC Grant agreement n° [648030] awarded to PB and by the Lundbeck Foundation BRAINSTRUC initiative (R155-2015-2666). The CBS is a member of France-BioImaging (FBI) and the French Infrastructure for Integrated Structural Biology (FRISBI), 2 national infrastructures supported by the French National Research Agency (ANR-10-INBS-04-01 and ANR-10-INBS-05, respectively).

