## Supplementary Information for "WASCO: A Wasserstein-based statistical tool to compare conformational ensembles of intrinsically disordered proteins"

### S1 Methodology details

#### S1.1 Building a residue-specific reference frame

##### *S1.1.1 Reference frame definition*

We seek to define a reference frame that determines the global pose (position and orientation) of a given residue and that allows to describe the relative pose of other residues along the sequence. As we want this reference system to be universally defined (independently of the residue identity), we first define a virtual atom $\widetilde{C_\beta}$, which exists also for glycines. The position of $\widetilde{C_\beta}$ is an estimate of the position of the true $C_\beta$ when it exists, but it is defined for every residue using only the atoms that are always present. Its definition allows the construction of a universal frame that locally represents the geometry of the backbone.

Let $\overrightarrow{C}$ and $\overrightarrow{N}$ be the vectors going from $C_\alpha$ to $C$ and $N$ atoms, respectively. If a $C_\beta$ atom is present, let $\overrightarrow{C_\beta}$ denote the vector going from $C_\alpha$ to $C_\beta$. In such case, $\overrightarrow{C_\beta}$ can be determined using the vectors $\overrightarrow{C}$, $\overrightarrow{N}$ and $\overrightarrow{C} \times \overrightarrow{N}$ together with their angles with respect to $\overrightarrow{C_\beta}$, denoted $\theta_C$, $\theta_N$ and $\theta_{CN}$ respectively. See Figure S1a for an illustration. This can be done by solving the following linear system, whose unknown variables are the three coordinates of $C_\beta$.

$$\begin{cases} \|\overrightarrow{N}\| \, \|\overrightarrow{C_\beta}\| \, \cos\theta_N = \overrightarrow{N} \cdot \overrightarrow{C_\beta} \\ \|\overrightarrow{C}\| \, \|\overrightarrow{C_\beta}\| \, \cos\theta_C = \overrightarrow{C} \cdot \overrightarrow{C_\beta} \\ \|\overrightarrow{C} \times \overrightarrow{N}\| \, \|\overrightarrow{C_\beta}\| \, \cos\theta_{CN} = (\overrightarrow{C} \times \overrightarrow{N}) \cdot \overrightarrow{C_\beta}. \end{cases} \tag{1}$$

To define a *universal* $C_\beta$, denoted $\widetilde{C_\beta}$, we will estimate fixed values for $\theta_N$, $\theta_C$ and $\theta_{CN}$ from all non-glycine residues of a set of protein structures and *define* the $\widetilde{C_\beta}$ coordinates as the solution of (1), independently of the residue identity. Details on angles estimation are given in the following section. Consequently, for a given residue, the virtual atom $\widetilde{C_\beta}$ is determined from the coordinates of its $C_\alpha$, $N$ and $C$ atoms. This allow us to define a reference system at each sequence position through the following three vectors, where $\overrightarrow{CN} = \overrightarrow{N} - \overrightarrow{C}$.

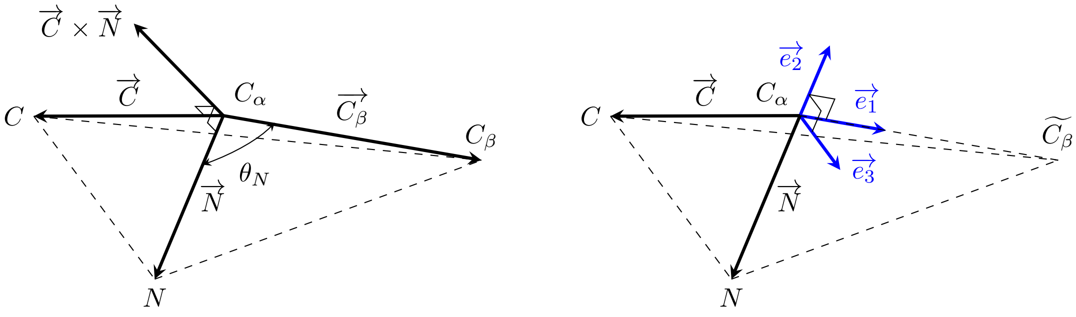


(a) (b)

Figure S1: (a) Illustration of vectors and angles involved in the construction of the residue-specific reference frame. The vector $\overrightarrow{C_\beta}$ can be determined from vectors $\overrightarrow{C}$ and $\overrightarrow{N}$ together with the angles $\theta_N$ (the only depicted for simplicity), $\theta_C$ and $\theta_{CN}$. (b) The three vectors $\{\overrightarrow{e_1}, \overrightarrow{e_2}, \overrightarrow{e_3}\}$ defining the reference frame, built from the virtual atom $\widetilde{C_\beta}$ and vectors $\overrightarrow{C}$ and $\overrightarrow{N}$.

$$\begin{cases} \overrightarrow{e_1} = \overrightarrow{\widetilde{C_\beta}}/\|\overrightarrow{\widetilde{C}_\beta}\| \\ \overrightarrow{e_2} = \overrightarrow{CN}/\|\overrightarrow{CN}\| \times \overrightarrow{e_1} \\ \overrightarrow{e_3} = \overrightarrow{e_1} \times \overrightarrow{e_2}. \end{cases} \tag{2}$$

Once the reference system of the $i$-th residue, denoted $\mathcal{F}_i = \{\overrightarrow{e_{1,i}}, \overrightarrow{e_{2,i}}, \overrightarrow{e_{3,i}}\}$, has been built, its origin will be placed at the $C_\beta$ atom when it exists, or at the $C_\alpha$ otherwise. This allows the computation of relative positions and distances with respect to $C_\beta$ atoms for all non-glycine residues.

#### *S1.1.2 Estimation of $\theta_C$, $\theta_N$ and $\theta_{CN}$*

We estimated three fixed values for $\theta_C$, $\theta_N$ and $\theta_{CN}$, to be replaced in the linear system (1). After that, the vector $\overrightarrow{\widetilde{C_\beta}}$ is determined for each residue along the sequence by solving (1) after plugging in the corresponding coordinates of $C_\alpha$, $C$ and $N$ atoms. As we mentioned in Section S1.1, this allows the definition of a residue-specific reference frame, built independently of the residue identity.

To estimate the three angles, we used a set of 15177 experimentally-determined high-resolution structures of protein domains extracted from the SCOPe 2.07 release [1]. For each structure, $\theta_C$, $\theta_N$ and $\theta_{CN}$ were computed and stored for every non-glycine residue. The three corresponding histograms, together with a kernel density estimate, are presented in Figure S2, for all residue types. The residue-specific counterparts of Figure S2 did not show important fluctuations from the overall densities. Therefore, for simplicity, we did not estimate three angles per residue type, but three universal values.

The three distributions of Figure S2 show that all the angle distributions are strongly concentrated around their kernel density maximum. Consequently, these values were chosen as an estimate of $\theta_C$, $\theta_N$ and $\theta_{CN}$. Due to the symmetry of the empirical distributions, choosing the mean would provide similar estimates. Figure S2 depicts the theoretical angle values under the hypothesis that $C$, $N$, $C_\beta$ and $H$ (when present) are the vertices of a regular tetrahedron, with $C_\alpha$ as its centroid. One could think of using these values as estimates, but the deviation from the experimental value of $\theta_{CN}$ is too high, showing how the fluctuations from the regular polyhedron are not homogeneous along its faces.

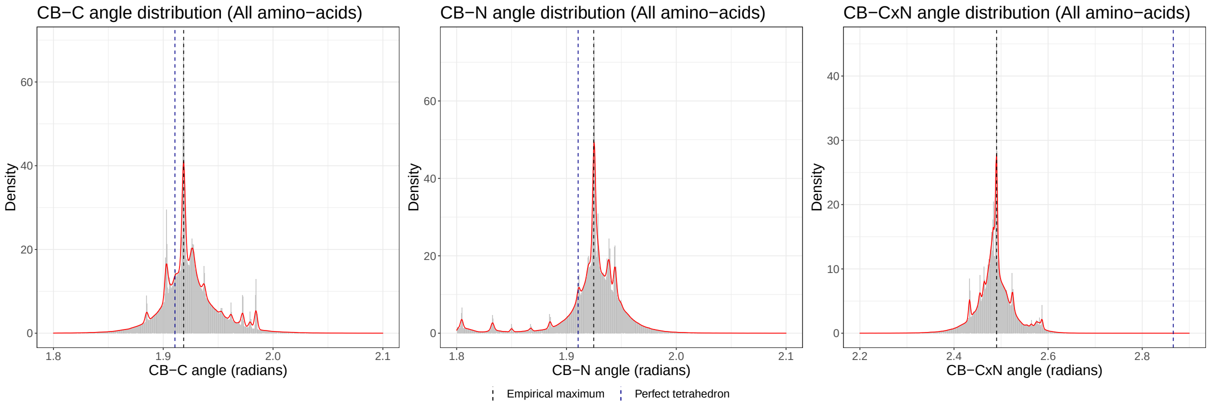


Figure S2: From left to right: empirical distributions of $\theta_C$, $\theta_N$ and $\theta_{CN}$ respectively, extracted from a set of 15177 protein structures, considering all non-glycine residues. The red line corresponds to a kernel density estimate, whose maximum (vertical black dashed line) was used as angle estimate. The blue dashed line depicts the theoretical value of each angle under the hypothesis that the four atoms bound to the $C_\alpha$ form a regular tetrahedron.

#### S1.2 Practical computation of Wasserstein distance

In this work, we consider the 2-Wasserstein distance and refer to it simply as the Wasserstein distance. The Wasserstein distance can be easily computed from a pair of samples drawn from the corresponding probability distributions. However, a major drawback of the algorithms that compute the Wasserstein distance is their inability to handle large datasets ($\gtrsim 10^3$ points). The current implementations in Python [2] or R [3] only admit datasets with $\lesssim 5 \cdot 10^3$ points, which is usually not enough for conformational ensembles. To the best of our knowledge, there are no existing algorithms that solve an OT problem for large sample sizes and which are easily implementable, considerably fast (which, in our case, is essential due to the large number of Wasserstein distances to compute) and which accept non-euclidean ground distances (like the distance in the torus).

Here, we propose an approximation method to "simplify" the input empirical distributions and compute the Wasserstein distance from a pair of smaller samples sizes. The efficiency of this approach in terms of error is illustrated via simulations on real protein data, but we provide no theoretical bounds. The proposed algorithm consists on clustering the original distribution and define its clustered version as a discrete probability distribution supported on the set of clusters whose mass is given by the proportion of points assigned to each. Then, the Wasserstein distance is computed between the pair of clustered distributions, whose samples have admissible sizes. The method is implemented for both local and global structural descriptors, which are empirical probability distributions supported on $\mathbb{T}^2$ and $\mathbb{R}^3$ respectively.

The accuracy in terms of relative and mean-square error is presented in Figure S3. Note that the approximation algorithm has a considerably better performance when implemented for local structural descriptors, which was expected due to the boundedness of the corresponding ground space. Accuracy in $\mathbb{R}^3$ is slightly worse, as cloud points representing the relative position of residues are in general more disperse, and therefore the clustered distribution needs a larger number of centroids to better capture its variability. Nevertheless, we observe that in both cases the error estimates for a proportion of $\sim 10\%$ of clusters with respect to the entire dataset size (the proportion we will be using in practice) are acceptable for our practical purposes. To enrich interpretation, we performed the same accuracy analysis but computing Wasserstein distance between subsamples uniformly drawn from the corresponding datasets. As it is shown in Figure S3, the effect of clustering considerably improves the quality of the approximation.

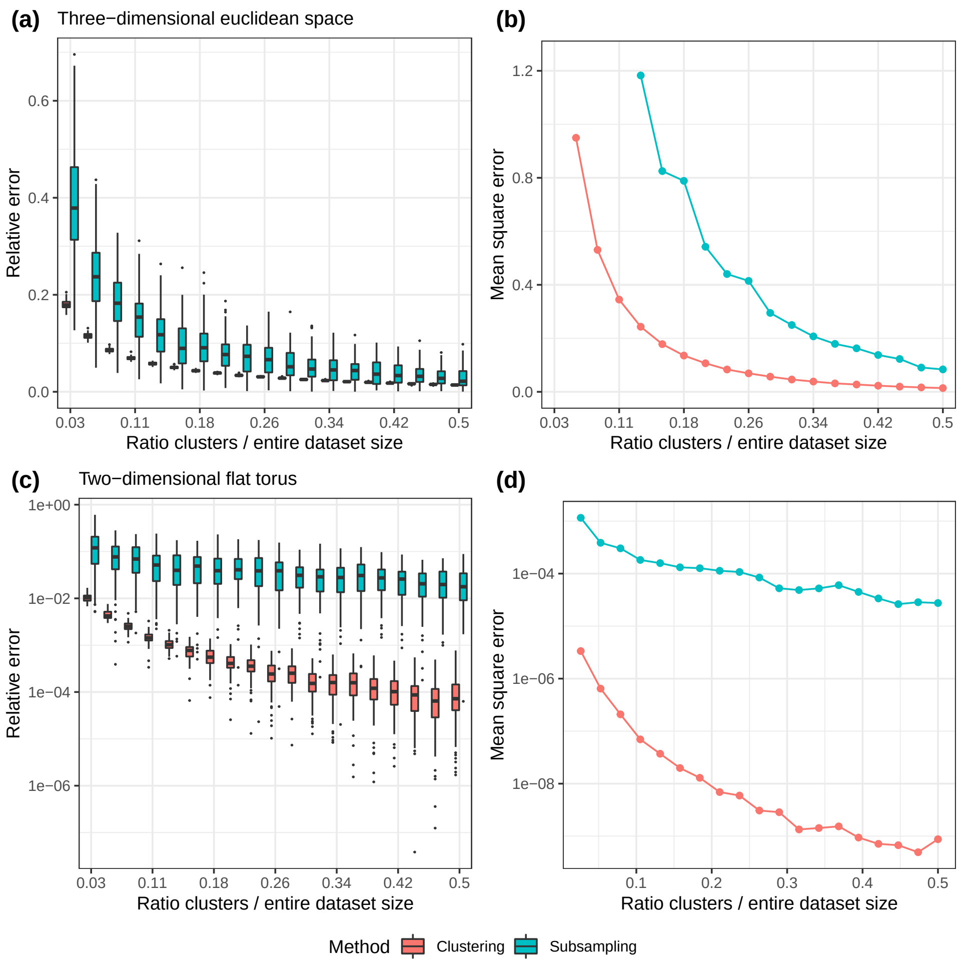


Figure S3: From left to right (columns): relative and mean square error estimates of the Wasserstein distance between the clustered distribution as an estimate of the Wasserstein distance between the original datasets. In abscissas, the proportion of the number of clusters with respect to the entire dataset size. The first row (a,b) corresponds to samples drawn from local structural descriptors (dihedral angles) and the second (c,d) to samples drawn from global structural descriptors (pairwise relative positions of residues).

### S2 Supplementary figures

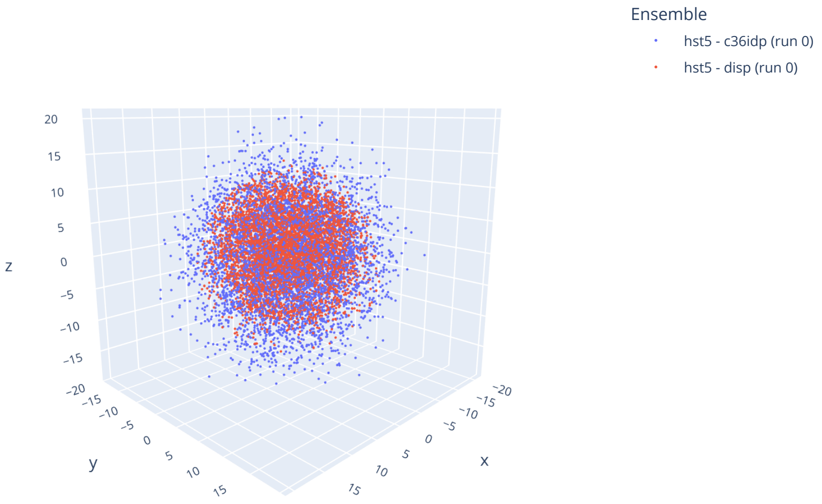


Figure S4: Two samples of $\overrightarrow{R}_{3,10}$ corresponding to a pair of ensembles of Hst5 simulated with force-fields CHARMM36IDPSFF (c36idp) and AMBER ff99SB-disp (disp). Each sample is represented by a point cloud in the three-dimensional euclidean space.
